# Detection of Envelope Glycoprotein Assembly from Old-World Hantaviruses in the Golgi Apparatus of Living Cells

**DOI:** 10.1101/2020.06.01.127639

**Authors:** R. A. Petazzi, A. A. Koikkarah, N.D. Tischler, S. Chiantia

**Affiliations:** University of Potsdam, Institute of Biochemistry and Biology, Karl-Liebknecht-Str. 24-25, 14476 Potsdam, Germany; Fundación Ciencia & Vida, Laboratorio de Virología Molecular, Av. Zañartu 1482, Santiago, Chile; Universidad San Sebastián, Santiago, Chile

**Keywords:** fluorescence fluctuation microscopy, number and brightness, virus assembly, fluorescence correlation spectroscopy, protein-protein interaction

## Abstract

Hantaviruses are emerging pathogens that occasionally cause deadly outbreaks in the human population. While the structure of the viral envelope has been characterized with high precision, the protein-protein interactions leading to the formation of new virions in infected cells are not fully understood yet. In this work, we use quantitative fluorescence microscopy (i.e. Number&Brightness analysis and fluorescence fluctuation spectroscopy) to quantify the interactions that lead to oligomeric spike complex formation in the physiological context of living cells. To this aim, we have analyzed proteins from Puumala and Hantaan orthohantaviruses in several cellular models. For the first time, we quantified the oligomerization state of each protein in relation to its subcellular localization, concentration and the concentration of its interaction partner. Our results indicate that when expressed separately, both glycoproteins can form homo-multimers in a concentration-dependent manner. Fluorescence fluctuation analysis was applied to prove that Gc:Gc contacts observed on virions are also relevant for Gc-Gc interactions in living cells, in the absence of Gn. Furthermore, we proved that the membrane-distal lobes of Gn are not necessary for Gn homo-multimerization. In cells co-expressing both glycoproteins, we observe clear indication of Gn-Gc interactions and the formation of protein complexes with different sizes, while using various labelling schemes to minimize the influence of the fluorescent tags. Our data are compatible with an assembly model according to which hantavirus spikes are formed via the assembly of Gn-Gc hetero-dimers. Furthermore, our results indicate the interconnection of large Gn-Gc hetero-multimers in the Golgi apparatus. Such large glycoprotein multimers may be identified as multiple interacting viral spikes and provide a possible first evidence for the initial assembly steps of the viral envelope, within this organelle, directly in living cells.

## Introduction

Hantaviruses (HV) are single-stranded, negative sense RNA viruses belonging to the *Hantaviridae* family of the order Bunyavirales. While usually infecting rodents, insectivorous mammals or bats (1), these emerging pathogens can occasionally cause deadly outbreaks in the human population (2). Their segmented genome encodes at least a nucleocapsid protein, a RNA-dependent RNA polymerase and a glycoprotein precursor (GPC) (3). The assembly of HV particles in infected cells is a complex process that relies on the precise spatial and temporal organization of several viral proteins. The specific molecular mechanisms driving protein-protein interactions in this context are not fully characterized yet.

While negative-stranded RNA viruses typically assemble and bud at the plasma membrane (4, 5), Bunyavirales are mostly characterized by intracellular maturation (6). In all HVs, two glycoproteins (GPs) form surface spikes and are derived from the GPC encoded in the M segment of the tripartite genome. After, The N-terminal Gn and C-terminal Gc have been suggested to be cleaved co-translational by cellular signal peptidases (7). The interplay between Gn and Gc has a fundamental importance in HV assembly and budding (8, 9).

Early studies that aimed to localize singularly expressed Gn and Gc showed variations for different members of the Bunyavirales order (10, 11). However, co-expression of both HV GPs results, in general, in their translocation to the Golgi apparatus (GA) (12-16). This holds true also for recombinant proteins that are not derived from a common precursor (17).

On the other hand, not much is known about where and how Gn-Gc hetero-complexes are initially assembled in an infected cell. It is therefore of great interest to address the role of protein-protein interactions involved in the maturation of the HV spike complex. Cryo-EM studies on the surface of Tula virus (TULV) indicated that such a complex is formed by square-shaped Gn-Gc hetero-octamers composed of four Gn and Gc molecules (18, 19). The presence of Gn-Gc heteromultimers has also been confirmed by biochemical assays of virion extracts (20). The fitting of the Gn ectodomain crystal structure of PUUV into the spike density of a cryo-EM tomography reconstruction of TULV revealed that Gn forms the membrane distal tetrameric lobes with Gc occupying the volume underneath (21). The design of cysteine mutations at the two-fold axis of a crystallographic Gc dimer (22) further proved that Gc:Gc contacts connect adjacent spikes and additional residue substitutions showed that the interactions at the inter-spike Gc:Gc interface are crucial for virus particle assembly (23).

A quantitative assessment of Gn-Gc interactions leading to the formation of spike complexes in a more physiological environment within a living cell is still lacking. According to the model proposed by Hepojoki *et al*. (24), the assembly of HV spikes might follow two pathways. In one case (referred to as “assembly model 1” in what follows), homotypic interactions between Gn monomers lead first to the formation of a tetrameric core to which Gc proteins subsequently bind to form larger hetero-complexes. In the second case (i.e. “assembly model 2”), the formation of a Gn-Gc heterodimer is the first step in the formation of the final spike complex (via interactions between four Gn-Gc heterodimers).

While the above-mentioned biochemical and structural characterizations of virion surface provide information about completely formed spike complexes, alternative approaches are needed to clarify how protein-protein interactions occur in different intra-cellular compartments, leading to the spike and inter-spike assembly.

Here, we apply Number and Brightness (N&B) analysis to quantify the homo- and hetero-oligomerization of HV Gn and Gc, directly in living cells. Previously, we have used this method to demonstrate that Gn possesses a significantly higher tendency to oligomerize, compared to Gc. This approach was based on the bulk analysis of several cells, in a restricted protein concentration range, and required some assumptions on the actual presence of specific multimeric states (e.g. Gn tetramers) (16).

In this new work, we provide a complete quantitative characterization of HV GPs homo-multimerization, over a large concentration range, for two different virus strains and in different cell models. Also, we investigate the hetero-interactions between Gn and Gc using a novel quantitative fluorescence assay based on bidirectional expression vectors, allowing us to monitor the oligomerization of one protein as a function of the concentration of the other. Finally, we determine the most probable of the proposed assembly models and provide the first experimental evidence of HV assembly directly in the GA of living cells.

## Materials and Methods

### Cloning and generation of chimeric proteins

Unless otherwise specified, glycoprotein gene sequences refer to the PUUV strain. Original plasmids of signal peptide (SP)-mEYFP-Gn and SP-mEYFP-Gc were previously described (16). Plasmids encoding Gn and Gc preceded by three HA tags (HAtag-Gn, HAtag-Gc) were a gift from A. Herrmann (Humboldt Universität zu Berlin). A monomeric variant of EGFP, containing the A206K mutation (25), was PCR amplified from a mEGFP-C1 vector and cloned into these vectors to substitute mEYFP and generate SP-mEGFP-Gn and SP-mEGFP-Gc respectively. Vectors for the expression of N-terminally mEGFP-tagged Gn and Gc mutants were synthesized by Twist Bioscience (San Francisco, CA, USA): a Gn variant comprising of amino acids 387-658 and lacking part of the ectodomain (ΔGn) and Gc variants with point mutations at the Gc:Gc interface (H303C Gc, H303E Gc). ΔGn contains a flexible linker between mEGFP and the truncated Gn (EGKSSGSGSESKST) (26, 27). A plasmid for bi-directional expression was a gift from K. Arndt (Universität Potsdam). Promoters TTC31 and CCDC142 allowed simultaneous expression of encoded genes (28). mEGFP and mCherry2 were PCR-amplified and cloned into the two expression cassettes flanked by restriction sites BamHI/EcoRI and SacI/KpnI respectively, to obtain mEGFP⟵⟶mCherry2. A construct with mCherry2 and mEGFP cloned into SacI/KpnI and BamHI/EcoRI restriction sites, respectively, was also produced (mCherry2⟵⟶mEGFP). Unlabeled GP sequence (SP-Gn and SP-Gc) were PCR-amplified and separately cloned into the BamHI/EcoRI cassette to produce SP-Gn⟵⟶mCherry2 and SP-Gc⟵⟶mCherry2. A plasmid encoding the whole Hantaan virus (HTNV) GPC was a gift from Dr. S. Weiss (Charité, Berlin). The signal peptide was introduced at the N-terminus of mEGFP-C1 and Gn and Gc were independently amplified and cloned into this vector to obtain HTNV SP-mEGFP-Gn and SP-mEGFP-Gc. Golgi-mTurquoise was a gift from Dorus Gadella (Addgene plasmid # 36205) and has been described before (16). ER-mCherry, Golgi-mCherry and MyrPalm-mEGFP were previously described in (16). ER-mEGFP and Golgi-mEGFP were obtained by PCR amplification of mEGFP from mEGFP-C1 and subtitution of the FP via digestion/ligation at the restriction sites AgeI/BsrgI. FastDigest enzymes (Thermo Scientific, Waltham, MA, USA) were used for all restriction reactions. Ligations were performed with T4 DNA Ligase (Thermo Scientific) at 16° overnight. All cloning were tested by Sanger sequencing of expression plasmids.

### Cell culture and transfection

Chinese hamster ovary cells (CHO-K1), adenocarcinomic human alveolar basal epithelial cells (A549), human embryo kidney cells (HEK-293T) and VeroE6 cells were maintained in Dulbecco’s modified Eagle’s medium containing 10% fetal bovine serum, 100 U/mL penicillin, 0.1 mg/mL streptomycin and 4 mM L-Glutamine. Cells were passaged every 3–5 days, no more than 15 times. All solutions, buffers and media used for cell culture were purchased from PAN-Biotech (Aidenbach, Germany). 3-6 · 10^5^ cells were plated on glass-bottom 35-mm-diameter plates (CellVis, Mountain View, CA or MatTek Corp., Ashland, MA) 48 hours before experiments. Fusion protein expression plasmids were transfected into (70-90% confluent) cells using Turbofect™ (Thermo Scientific) according to the manufacturer’s protocol, 20-24 hours prior to experiments. VeroE6 cells were transfected with Lipofectamine™ 3000 (Thermo Scientific) instead, since they are not susceptible to transfection by Turbofect™.

### Confocal fluorescence microscopy

Confocal fluorescence images were obtained with a 40× water immersion objective (NA 1.2) at 21°C on a Zeiss LSM780 (Carl Zeiss, Oberkochen, Germany) confocal microscope with a frame size of 512 × 512 pixels. mTurquoise was excited at 405 nm using a laser diode and observed in 475–490 nm detection range. mEGFP was excited at 488 nm using a CW Argon laser (Lasos, Jena, Germany) and detected in the range of 498–606 nm. mCherry2 was excited at 561 nm using a laser diode and observed in 578-695 nm detection range. Images with more than one fluorescent species were recorded sequentially to minimize signal cross-talk.

For intracellular immunofluorescence staining, cells were washed three times with phosphate-buffered saline with calcium and magnesium (PBS, PAN-Biotech) and fixed with 3.7% paraformaldehyde (Sigma-Aldrich, Munich, Germany) for 25 min at room temperature. Afterwards, the cells were washed three times with PBS before permeabilization with 0.2% Triton X-100 (Sigma-Aldrich) and 0.2% bovine serum albumin (PAN-Biotech) for 20 min. After three more washing steps, cells were incubated with anti-HAtag rabbit primary antibody (Abcam, Berlin, Germany, ab13511) for 1 h. This procedure was repeated for the Alexa488-conjugated secondary antibody (goat anti rabbit IgG, Thermo Scientific) and the immunostaining was concluded with three washing steps.

### Fluorescence Correlation Spectroscopy

Point FCS measurements (29) were performed for 120 s on a Zeiss LSM780 microscope and recorded using the Zen Black software. mCherry2 was excited at 561 nm using a laser diode and observed in 578-695 nm detection range. Laser powers were adjusted to keep photobleaching below 20%. Typical values were ∼6 µW (561 nm). The size of the confocal pinhole was set to 1 airy unit. Intensity time series were analyzed with the Zen software to obtain the auto-correlation function (ACF):

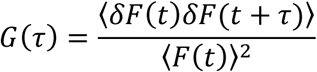

where *δF*(*t*) = *F*(*t*) − ⟨*F*(*t*)⟩. The ACF was fitted as described in (30). The inverse of the amplitude G(0) indicates the number of particles in the effective confocal volume (29). To calibrate the confocal volume, pFCS measurements with Alexa Fluor® 488 or Rhodamine B dissolved in water at 50 nM were performed at the same laser power. The structure parameter S (defined as the ratio between the vertical and lateral dimension of the theoretical confocal ellipsoid) was fixed to the value determined in the calibration measurement (typically around 4 to 8).

### Number and Brightness

Number and Brightness analysis (N&B) was performed as previously described (16, 31). Briefly, 3-6 · 10^5^ cells were plated onto 35 mm glass-bottom dishes (CellVis, Mountain View, CA or MatTek Corp., Ashland, MA) 48 h prior to the experiment and transfected as described above. Confocal images were acquired using a Zeiss LSM780 microscope. The 488 nm excitation from a CW Argon laser was focused with a 40× UPLS Apochromat 1.2 NA water objective into the sample. The fluorescence signal was collected by a Zeiss QUASAR multichannel GaAsP detector in a 498-606 nm range in photon counting mode. The laser power was set so that the photon count rate and bleaching remained below 1 MHz and ca. 20%, respectively (i.e. typical laser power values were ≤ 4 μW). Images of 128 × 128 pixels were acquired with pixel dimensions ∼ 400 nm and a pixel dwell time of 25-50 μs. Image time-stacks of 105 scans were collected using the Zeiss Black ZEN software. The intensity time-stacks data were analyzed using a self-written Matlab code (The MathWorks, Natick, MA, USA). The Matlab algorithm implements the equations from Digman *et al*. (32) for the specific case of photon-counting detectors, thus obtaining the molecular brightness and number as a function of pixel position. Regions of interest (e.g. ER, GA) were selected in cell images by applying a mask obtained from the intensity map of co-transfected ER-mCherry2 or Golgi-mCherry2.

To correct for photobleaching effects and minor cell movements, a boxcar-filter pixel-wise with 8-frames length was used, as previously described (16, 33). Saturation of detectors leading to artefactual reduction in brightness was avoided by excluding pixels in which photon-counting rates exceeded ∼ 1 MHz. Using this selection criterion, saturation induced brightness decrease was kept below 10 %. To further correct the detector response, we took into account the signal originating from a thin film containing immobilized fluorophores (31).

Finally the brightness values obtained from every single measurement were normalized to account for the monomer brightness *ε*_*m*_ and the fluorescence probability (*p*_*m*_), which determines the detectability of the tag (30). Briefly, the absolute brightness *ε*_*i*_ was compared to the brightness of the monomer and corrected using previously determined values for *p*_*m*_, thus obtaining an estimation of the multimerization state:

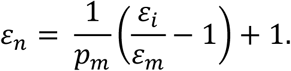

HV GPs are mostly found in the ER and GA (13, 14, 16, 34). Therefore, *ε*_*m*_ was determined for standard fluorescent proteins localized in the same subcellular compartments: ER-mEGFP (to be compared to Gn) and Golgi-mEGFP (to be compared to Gc). The actual multimeric state of these specific reference constructs was never quantified before. For this reason, the corresponding monomeric molecular brightness reference values were obtained by subjecting the specimen to several bleaching cycles (10-20s at a laser power 5-10 times higher than the measuring parameters) followed by N&B data acquisition. Since completely bleached multimers do not contribute to the overall brightness value, the lowest constant brightness measurement obtained indicates indeed the monomer reference brightness value (Fig. S1). We found that ER-mEGFP and Golgi-mEGFP form multimers (containing ca. 3-6 units) and that the monomeric brightness reference value can be calculated in a reproducible manner after the step-wise bleaching procedure (∼1 kHz/molecule). Furthermore, we observed that the monomer brightness values for the ER and GA markers were comparable to that of a membrane-associated monomeric fluorescent construct (i.e. MyrPalm-mEGFP) that is typically used in quantitative fluorescence microscopy experiments (16, 35). For this reason, we proceeded to use MyrPalm-mEGFP as reference for further measurements, thus avoiding the time-consuming step-wise bleaching calibration process.

The concentration N (in monomer units) was calculated by dividing the mean count rate in the ROI by the absolute brightness of the reference monomer:

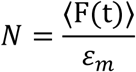

This corresponds to the number of monomers observed in the confocal volume (see previous paragraph). This number was then converted in concentration (i.e. monomers/µm^3^) using a value of 0.25 µm for the lateral size of the confocal volume. The structure parameter S of the confocal volume was the value determined in the calibration measurement, typically around 4 to 8. When plotting brightness vs. concentration data, segmented averages were calculated by dividing the whole concentration range in bins of variable width (containing ca. 1-4 points) and calculating mean values therein. Bins contained in general fewer points for high concentration values, i.e. in regions where the density of data points was lower. Unless differently stated, an empiric model was fitted to the data:

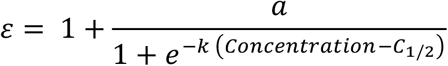

where *ε* is the normalized brightness and *a, k* and *C*_1/2_ are fitting parameters. *C*_1/2_ in particular is an indicator of the concentration at which half of the total protein is monomeric, in the simple approximation that only two oligomeric species are present (e.g. a mixture of 10 tetramers and 40 monomers).

Finally, in the case of mixtures of two different oligomeric states, the corresponding molar fractions n1 and n2 were determined by inverting the following equation:

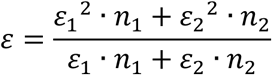

where *ε*_*i*_ indicates the multimerization of the i-th species and *ε* is the observed normalized brightness.

## Results

### Gn forms homo-tetramers in a concentration-dependent manner

In order to dissect the process of spike formation from GP monomers, we independently quantified the concentration-dependent multimerization of either Gn or Gc, at different protein concentrations, directly in living cells (36, 37). While local protein concentration cannot be actively controlled, it is possible to collect data from several cells randomly expressing different protein amounts. In order to quantify Gn (or Gc) multimerization, we performed N&B analysis. N&B is a fluorescence fluctuation-based method that allows to calculate the concentration and the molecular brightness of diffusing fluorescent molecules (32). The latter parameter indicates the amount of fluorescence signal detected for a single independent fluorescent molecular complex (e.g. a single protein tetramer) in the unit of time. The molecular brightness increases therefore if several monomers diffuse together (i.e. as a single entity) through the observation volume. By normalizing the measured value by the molecular brightness of a monomeric reference, it is possible to calculate the amount of fluorophores in the complex and, thus, its multimerization state (16, 30).

We analyzed mEGFP-tagged fluorescent constructs derived from two HV strains GPs in different cell line models (i.e. CHO-K1, HEK-293T, A549 and Vero E6 cells). In particular, the alveolar epithelial cells A549 were selected since “New World” HV strains can cause respiratory diseases (Hantavirus Pulmonary syndrome, HPS). Vero E6 cells are often used for the propagation and isolation of HVs. N-terminal labelling with mEGFP was preferred to C-terminal labelling based on previous results indicating the importance of GP C-terminus in protein localization (16). ER and GA markers – mCherry fusion proteins (FPs) bound to the respective retention peptides – were used to identify regions of interest (ROIs) and restrict the N&B analysis to specific cellular compartments.

In the case of PUUV GPs, SP-mEGFP-Gn was retained in the ER, while SP-mEGFP-Gc significantly partitioned in the GA (Fig. 1 A and C), as expected from previous studies (16) and controls with unlabeled GPs (Fig. S2). According to N&B analysis (Fig. 1 B), SP-mEGFP-Gn consistently showed a concentration-dependent oligomerization behavior, ranging from monomers to tetramers. Of interest, the measured brightness did not significantly exceed the values expected for tetramers, even at the highest observable concentrations.

**Figure 1:**
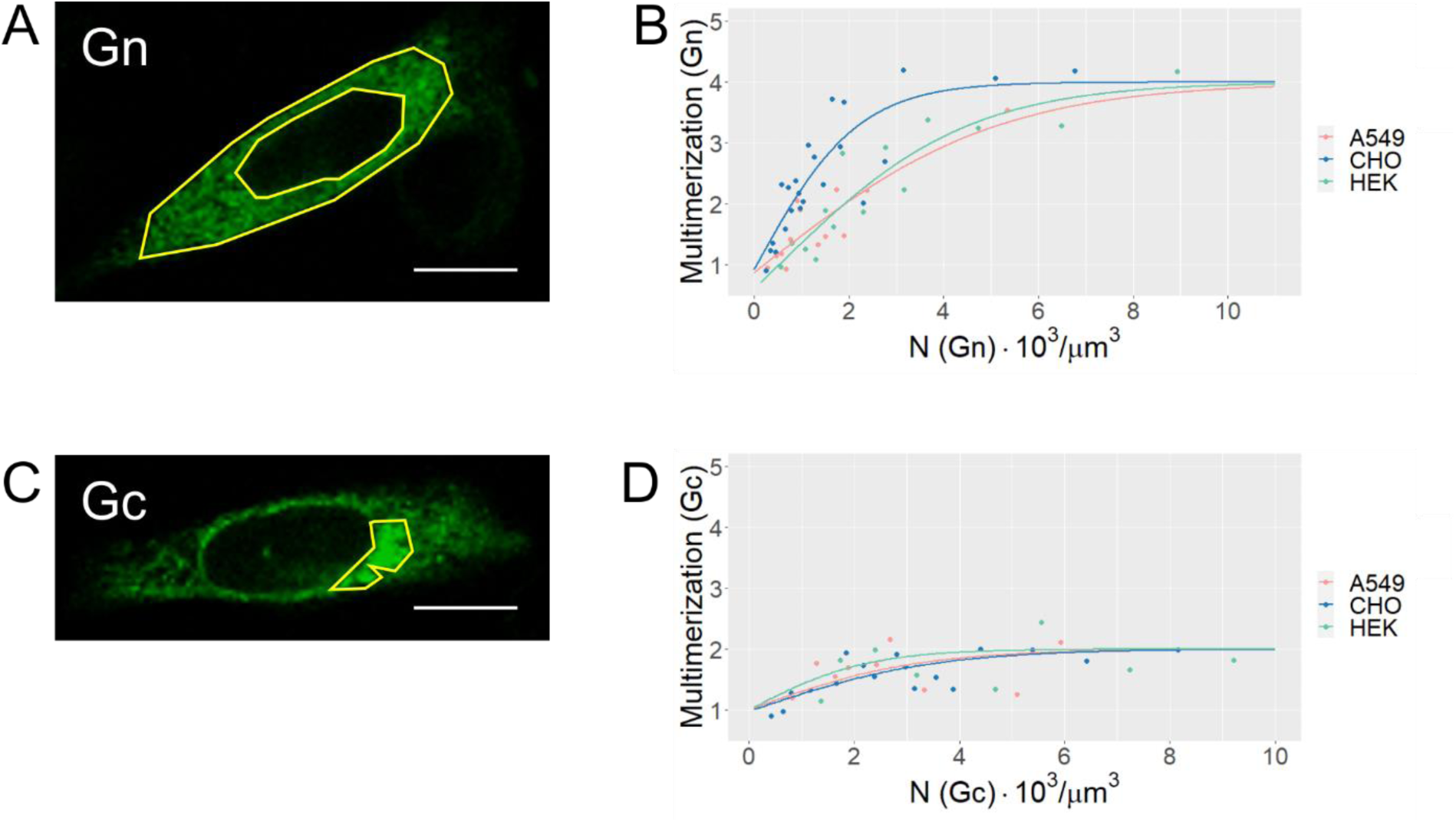
Concentration-dependent multimerization of PUUV Gc and Gn. Different cell lines expressed either SP-mEGFP-Gn (A, B) or SP-mEGFP-Gc (C, D). A, C) Representative fluorescence intensity maps of mEGFP-tagged GPs in CHO cells. Intensity maps of single cells were obtained as an average of 105 frames (128 × 128 pixels) collected over ∼2–3 min with a pixel dwell time of 25-50 μs. Examples or ROIs used for multimerization analysis are shown in yellow. White bars are 5 μm. B,D) Protein multimerization as a function of total protein concentration (in monomer units). N&B analysis was applied to a ROI in each cell (ER for Gn, Golgi for Gc) identified by an ER-mCherry or Golgi-mCherry marker. Multimerization values were obtained from molecular brightness normalization, as described in the Materials and Methods section. Each point is calculated from the average brightness extracted from ROIs in 2-4 cells with similar protein concentrations. Solid lines are a fit of an empirical model (see Materials and Methods) and are meant as guide to the eye. Each dataset is obtained from at least 5 independent experiments, for a total of approx. 50-70 examined cells.

The C½ concentration values (defined as the concentration at which half of the total protein is monomeric, in the simple approximation that only monomers and tetramers are present) range between ∼1·10^3^ (CHO) and ∼3·10^3^ (A549, HEK) monomers/µm^3^, for different cell lines.

On the other hand, SP-mEGFP-Gc multimerization only slightly correlates with its concentration in the GA (Fig. 1 D) or in the ER (Fig. S3). More in detail, SP-mEGFP-Gc oligomerization values ranged between those of monomers and dimers and did not exceed the value expected for dimers, even at the highest observable concentrations.

Analogous measurements were performed additionally in Vero E6 cells: although the expression of SP-mEGFP-Gn never reached concentrations high enough to observe a strong difference with SP-mEGFP-Gc or the clear formation of tetramers (Fig. S4, possibly due to sub-optimal transfection efficiency), GP multimerization might be compatible with that observed in other cell lines, at least at those low protein concentrations.

In order to assess whether HV GPs multimerization varies among different strains, Hantaan virus (HTNV) SP-mEGFP-Gn and SP-mEGFP-Gc were observed in CHO-K1 cells. Differently from PUUV GPs, both separately expressed proteins localize preferentially in the ER (Fig. 2 A and B). The quantification of the multimerization state via N&B analysis shown in Fig. 2 C indicates that SP-mEGFP-Gn forms monomers at low concentrations and tetramers at higher concentrations, similar to PUUV proteins. Also HTNV SP-mEGFP-Gc behaves similarly to its PUUV counterpart, displaying a multimerization state between monomers and dimers, over the explored concentration range.

**Figure 2:**
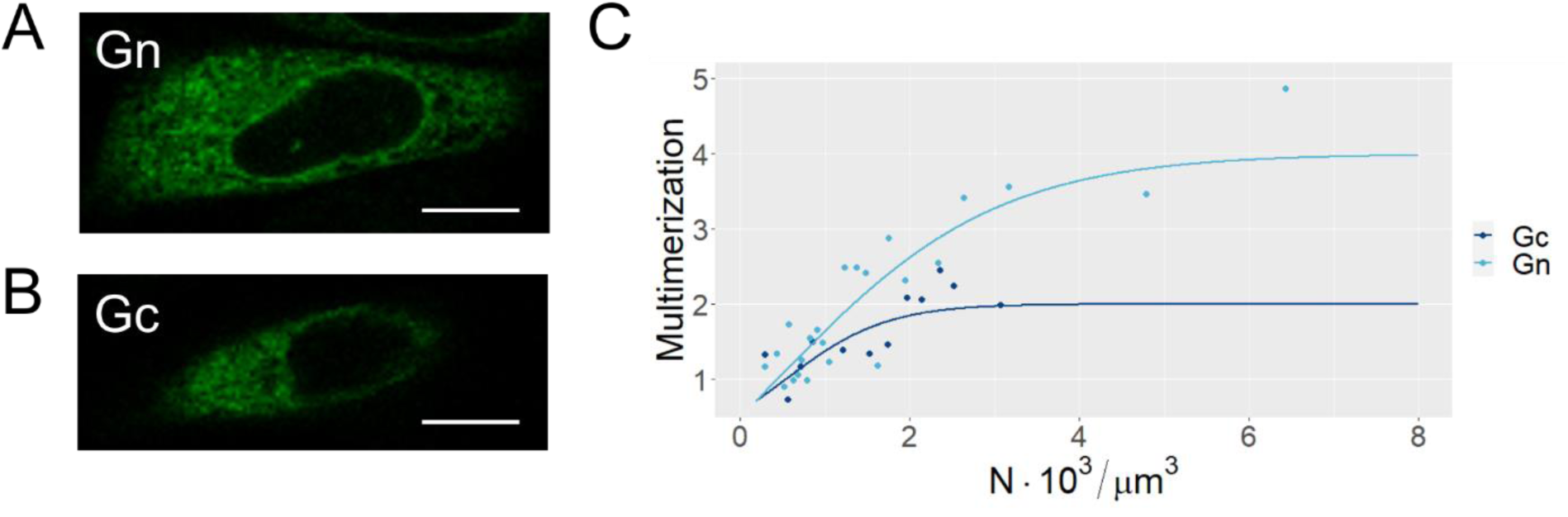
Concentration-dependent oligomerization of HTNV Gc and Gn. CHO-K1 cells were transfected with HTNV SP-mEGFP-Gn and SP-mEGFP-Gc vectors, respectively. A, B) Representative fluorescence intensity maps of mEGFP-tagged GPs in CHO cells. Intensity maps of single cells were obtained as an average of 100 frames (128 × 128 pixels) collected over ∼2–3 min with a pixel dwell time of 25-50 μs. White bars are 5 μm. C) Protein multimerization as a function of total protein concentration (in monomer units). N&B analysis was applied to a ROI in each cell identified by an ER-mCherry marker. Multimerization values were obtained from molecular brightness normalization, as described in the Materials and Methods section. Each point is calculated from the average brightness extracted from ROIs in 2-4 cells with similar protein concentrations. Solid lines are a fit to an empirical model (see Materials and Methods) and are meant as guide to the eye. Each dataset is obtained from at least 5 independent experiments, for a total of ca. 40-60 examined cells.

Altogether, these results indicate that, irrespective of specific cell line model or HV strain investigated, Gn can form oligomers (up to tetramers) when singularly expressed in cells. Gc, on the other hand, is found in monomeric and dimeric states at all investigated concentration levels.

### Gc:Gc interactions observed on the viral surface occur in cells also in the absence of Gn-Gc complexes

The Gc:Gc inter-spike contact region has been previously investigated by structure-guided biochemical assays of Andes HV-like particles to determine which residues are involved in stabilizing the interaction (23). In particular, cysteine substitutions of selected residues at the Gc:Gc interface showed a higher stability of Gn-Gc hetero-multimers. The introduction of repulsive interactions by residues with the same charge at the Gc:Gc interface, instead, led to a decrease in protein-protein interactions. Since the Gc:Gc interface region is highly conserved among different HV strains (23), we analyzed the same mutants (i.e., H303C and H303E) for the mEGFP-tagged PUUV Gc construct. The aim of this experiment was to verify whether Gc:Gc contacts relevant for inter-spike interactions play also a role in determining interactions between Gc monomers in living cells.

Expression in of Gc mutants in CHO-K1 cells indicated that both constructs are homogeneously distributed in the ER (Fig. 3, A and B) but were expressed with significantly different efficiencies (data not shown). We then selected a shared concentration range (i.e. 1-5·10^3^ monomers / μm^3^) for the three Gc constructs – wild type and the two mutants – and compared protein multimerization for these samples. Fig. 3 C shows that, compared to wt Gc, the H303E mutant displays a significantly lower multimerization. In the approximation that only Gc monomeric and dimeric assemblies are present (as suggested by the results reported in the previous paragraph), the observed average multimerization values correspond to a mixture of ca. 4:1 monomers:dimers for this mutant (cfr. approx. 3:2 monomers:dimers for the wt). On the other hand, H303C Gc mutants appear to be present almost exclusively in dimeric form, as expected from previous *in vitro* investigations of virus-like particles (23).

**Figure 3.**
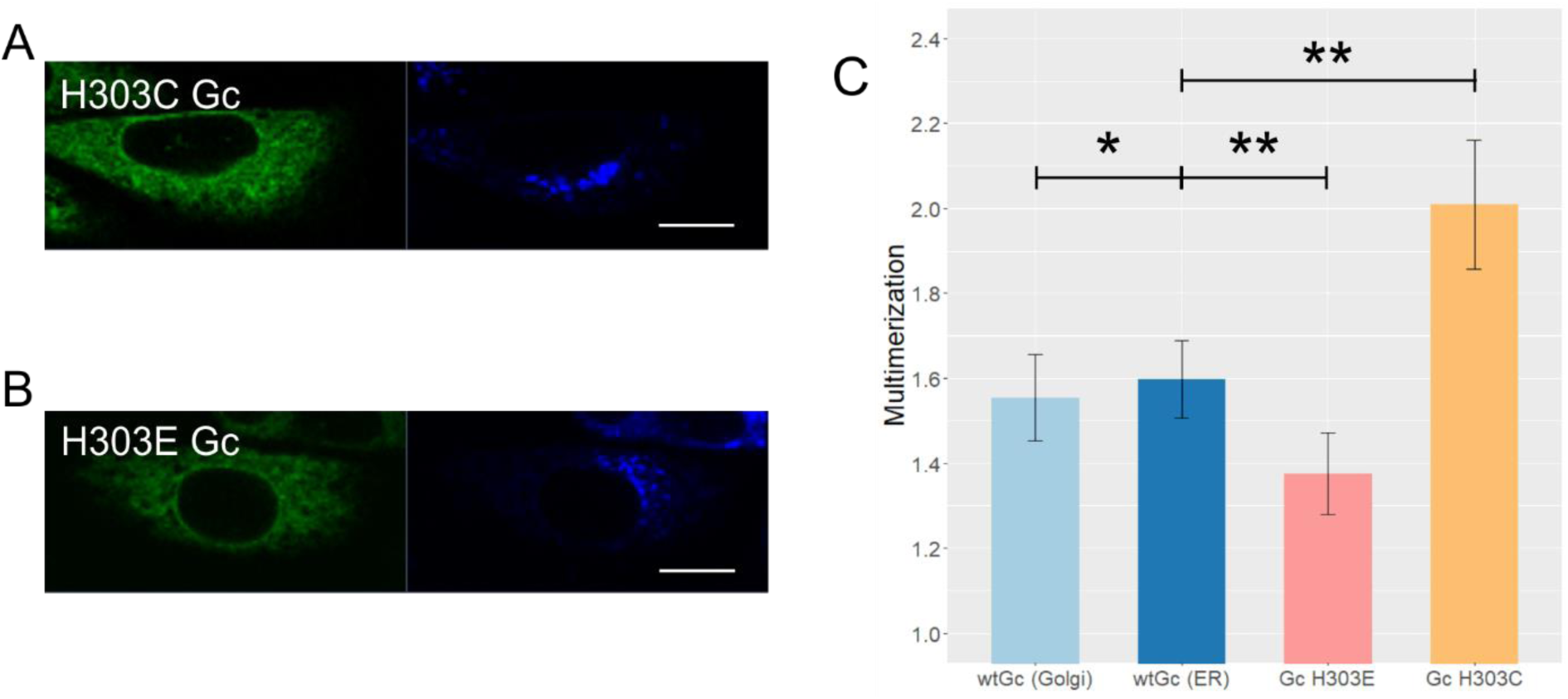
Brightness analysis of PUUV Gc mutants. CHO-K1 cells were transfected with SP-mEGFP-Gc(H303C), SP-mEGFP-Gc(H303E) or SP-mEGFP-Gc (wt) vectors. A, B) Representative fluorescence intensity images of PUUV Gc mutants H303C and H303E (left panels, in green) and Golgi-mTurquoise marker (right panels, in blue) co-expressed in CHO-K1 cells. Intensity images were obtained as an average of 105 frames (128 × 128 pixels) collected over ∼2–3 min with a pixel dwell time of 25-50 μs. White bars are 5 μm. C) Normalized multimerization values calculated via N&B analysis. In order to compare all the different constructs displaying varying expression efficiencies, multimerization values were selected in the shared range of 1-5·103 monomers / µm^3^. Bars indicate the median of at least 3 separate experiments and ca. 30 cells, error bars are calculated as the standard error of the mean. T-test significance is represented by asterisks: p > 0.5 (*), p < 0.05 (**).

These results indicate that Gc:Gc dimerization in our cellular model occurs through the same contacts, as on the viral surface between spikes. Also, N&B analysis can be successfully applied to detect subtle changes in Gc-Gc interactions (e.g. induced by point mutations) and such interactions do not appear to be significantly hindered by the presence of a fluorescent tag.

### The membrane-distal lobes of PUUV Gn are not necessary for tetramer formation

Based on the structural characterization of HV spikes, the central stalk of the Gn ectodomain (aa 388 to 484), Gn transmembrane region and endodomain may be responsible for homo-oligomerization (15, 21). Based on this criteria, we designed a Gn mutant lacking the membrane distal lobes of the ectodomain (21), thus obtaining SP-mEGFP-linker-Gn(387-658) (ΔGn). Protein localization was not influenced by the lack of the Gn lobes and the protein was still retained in the ER, similarly to wt SP-mEGFP-Gn (data not shown). As shown in Fig. 4, quantification of protein multimerization via N&B analysis indicates that ΔGn forms tetramers at high concentrations. The C½ value (ca. 5000 monomers / μm^3^) is significantly higher than the one observed for wt SP-mEGFP-Gn (ca. 1500 monomers / μm^3^, see Fig. 1 B and 4), implicating that tetramerization is favored in the case of the wt protein, although the Gn lobes are not strictly required.

**Figure 4:**
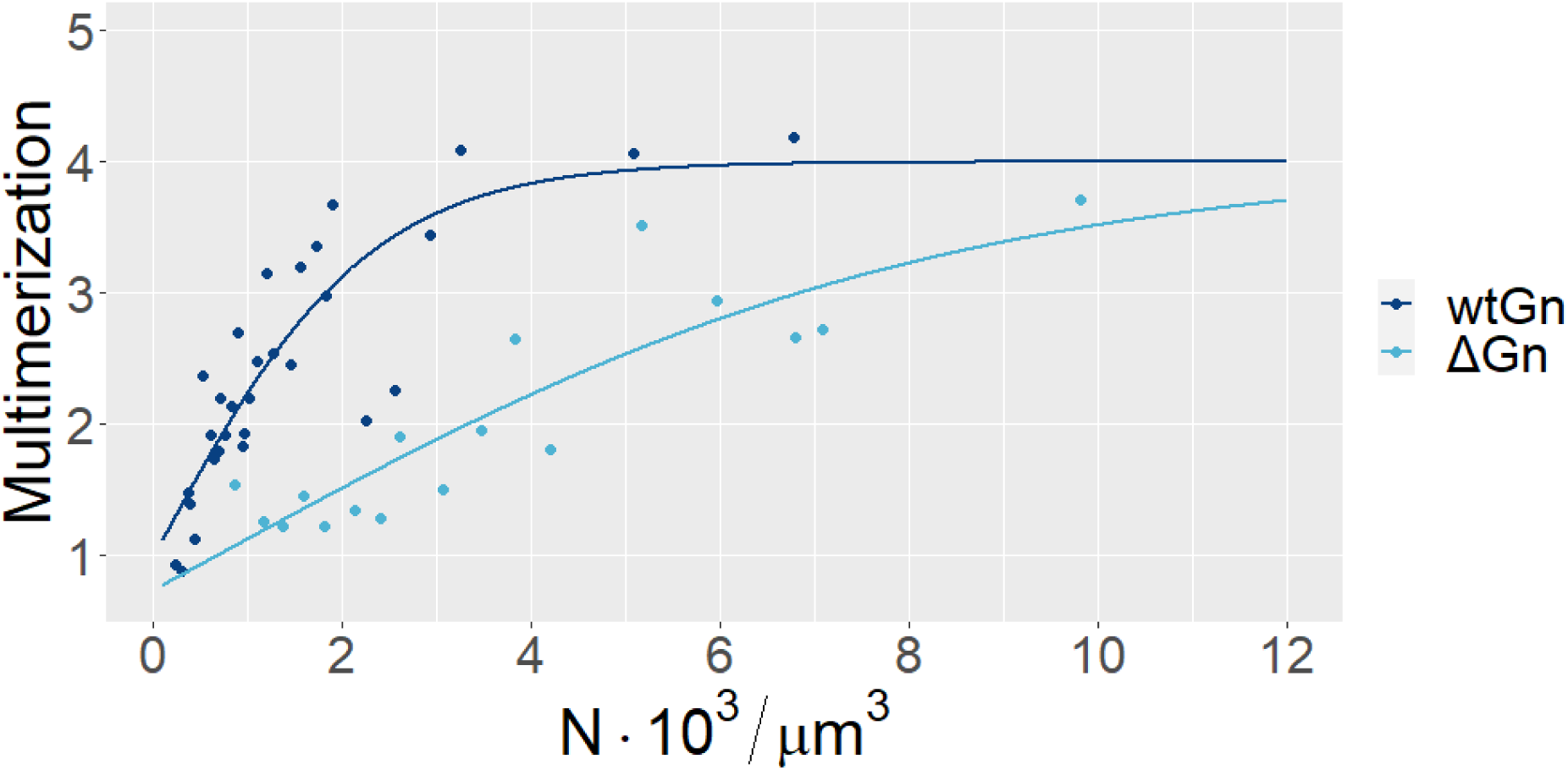
Comparative brightness analysis of PUUV Gn wt and ΔGn. CHO-K1 cells expressing ΔGn were analyzed. The mutant comprises amino acids 387-635 and is lacking part of the ectodomain. The graph shows protein multimerization as a function of total protein concentration (in monomer units), obtained via N&B analysis as described in the Materials and Methods section. The results for SP-mEGFP-Gn (wtGn) are shown as comparison. Each point is calculated from the average brightness extracted from ROIs in 2-4 cells with similar protein concentrations. Solid lines are a fit to an empirical model (see Materials and Methods) and are meant as guide to the eye. Each dataset is obtained from at least 5 independent experiments, for a total of ca. 50 examined cells.

### PUUV Gn forms high-order oligomers in the Golgi apparatus in the presence of Gc: 1) using fluorescent and untagged Gc constructs

We proceeded to study the interaction between Gn and Gc by co-expressing the two GPs in CHO-K1 cells and measuring their multimerization state via N&B analysis (Fig. 5). Co-expression of SP-mEGFP-Gn and SP-mCherry2-Gc caused an enrichment of both proteins to the GA, as previously observed (see Fig. S5 and (16)). The fluorescence signal (i.e. the protein concentration) in the ER was too low to obtain reproducible brightness values and, therefore, we quantified the multimerization of SP-mEGFP-Gn in the GA. Interestingly, Gn does not appear to form tetrameric assemblies (cfr. the case of SP-mEGFP-Gn in the absence of Gc, Fig. 1), even at relatively high concentrations (Fig. 5 E). This might be caused by the steric hindrance of FPs fused to the N-terminal of both GPs. In order to minimize this effect, we analyzed Gn multimerization also in cells co-expressing SP-mEGFP-Gn and unlabeled SP-Gc. While Gc is not detectable via fluorescence microscopy, we inferred its presence by the localization pattern of SP-mEGFP-Gn and limited the analysis to those cells with a clear partitioning of Gn in the GA. In these cases, SP-mEGFP-Gn was consistently found in higher-order multimers, up to octamers in average (Fig. S6).

**Figure 5:**
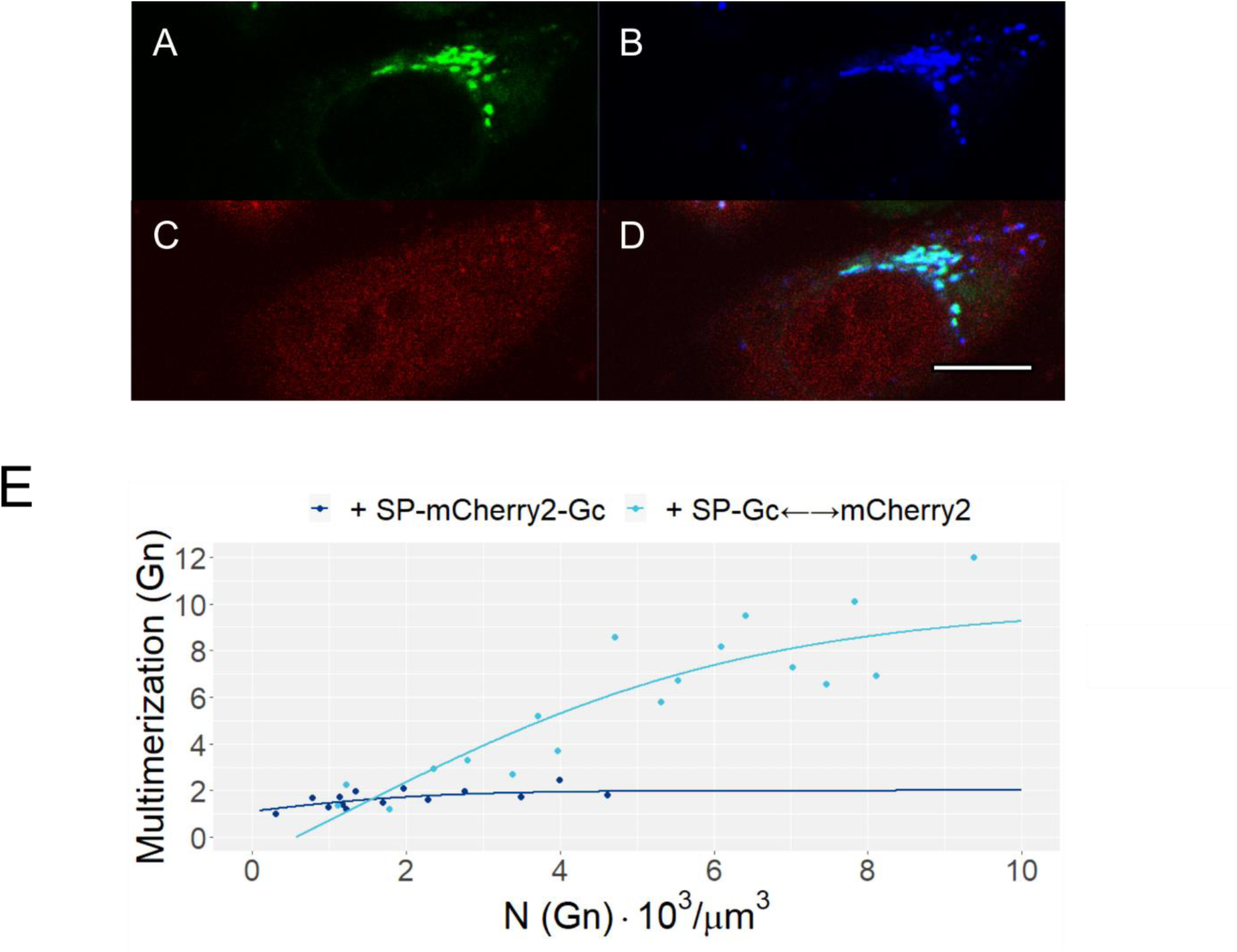
Concentration-dependent oligomerization of PUUV Gn in the presence of Gc. CHO-K1 cells expressed SP-mEGFP-Gn and either SP-mCherry2-Gc or SP-Gc (together with cytosolic mCherry2 within a bi-directional vector; indicated as SP-Gc⟵⟶mCherry2). A-D) Representative confocal microscopy image of CHO-K1 transfected with SP-mEGFP-Gn and SP-Gc⟵⟶mCherry2] vectors. Split view of SP-mEGFP-Gn (A), mGolgi-Turq marker (B), cytosolic mCherry2 (C) and merged channels (D). White bars are 5 μm. E) SP-mEGFP-Gn multimerization as a function of its total concentration (in monomer units), obtained via N&B analysis as described in the Materials and Methods section. The analysis was restricted to cells which showed significant expression of mCherry2. Each point is calculated from the average brightness extracted from ROIs in 2-3 cells with similar protein concentrations. Solid lines are a fit to an empirical model (see Materials and Methods) and are meant as guide to the eye. Each dataset is obtained from at least 4 separate experiments, for a total of ca. 40 cells.

### PUUV Gn forms high-order oligomers in the Golgi apparatus in the presence of Gc: 2) using bi-directional GP constructs

Transfecting cells with SP-mEGFP-Gn and an unlabeled SP-Gc construct allows an approximate characterization of Gn-Gc interactions. Although we focused on Gn localization in the GA as clue of the presence of Gc, this approach does not allow to confirm that both fluorescent Gn and unlabeled Gc are indeed present in every examined cell. We therefore adopted a novel alternative strategy that, additionally to solving the above-mentioned problem, allowed us to estimate the amount of unlabeled Gc. We transfected CHO-K1 cells with a vector for the expression of SP-mEGFP-Gn and, separately, a bi-directional vector for the concurrent expression of SP-Gc and cytosolic mCherry2 (indicated as SP-Gc⟵⟶mCherry2) (Fig. 5A-D). The measured concentration of mCherry2 can be then used to estimate the amount of unlabeled SP-Gc in the GA (Fig. S7). The concentration of SP-mEGFP-Gn outside the GA was too low to detect significant and reproducible protein multimerization. On the other hand, the quantification of SP-mEGFP-Gn oligomeric state as function of its concentration in the GA confirmed our initial estimations, showing higher average oligomerization states ranging up to roughly dodecamers (Fig. 5 E), far above the typical values observed for of SP-mEGFP-Gn in the absence of Gc. Of note, since the multimerization values do not reach a constant value at high concentrations, it must be assumed that of SP-mEGFP-Gn is, in general, present as a mixture of different multimeric species. In this case, N&B analysis provides simply a weighted average value for the multimerization state. Finally, we observed that SP-mEGFP-Gn multimerization state has only a weak correlation to the concentration of unlabeled SP-Gc (Fig. S8).

### PUUV Gc increases its multimerization in the GA in the presence of Gn

We proceeded to investigate the multimerization behavior of SP-mEGFP-Gc in the presence of SP-Gn⟵⟶mCherry2 (i.e. a non-labelled SP-Gn construct concurrently expressed with cytosolic mCherry2). While the intra-cellular localization of SP-mEGFP-Gc in the GA is not altered by the presence of the other GP (Fig. 6 A-D), its molecular brightness is significantly higher than that measured in the absence of Gn, with values ranging between two and four (Fig. 6 E). The apparent multimerization behavior of SP-mEGFP-Gc seems to be only weakly correlated to either its concentration (Fig. 6 E) or to SP-Gn concentration (Fig. S9).

**Figure 6:**
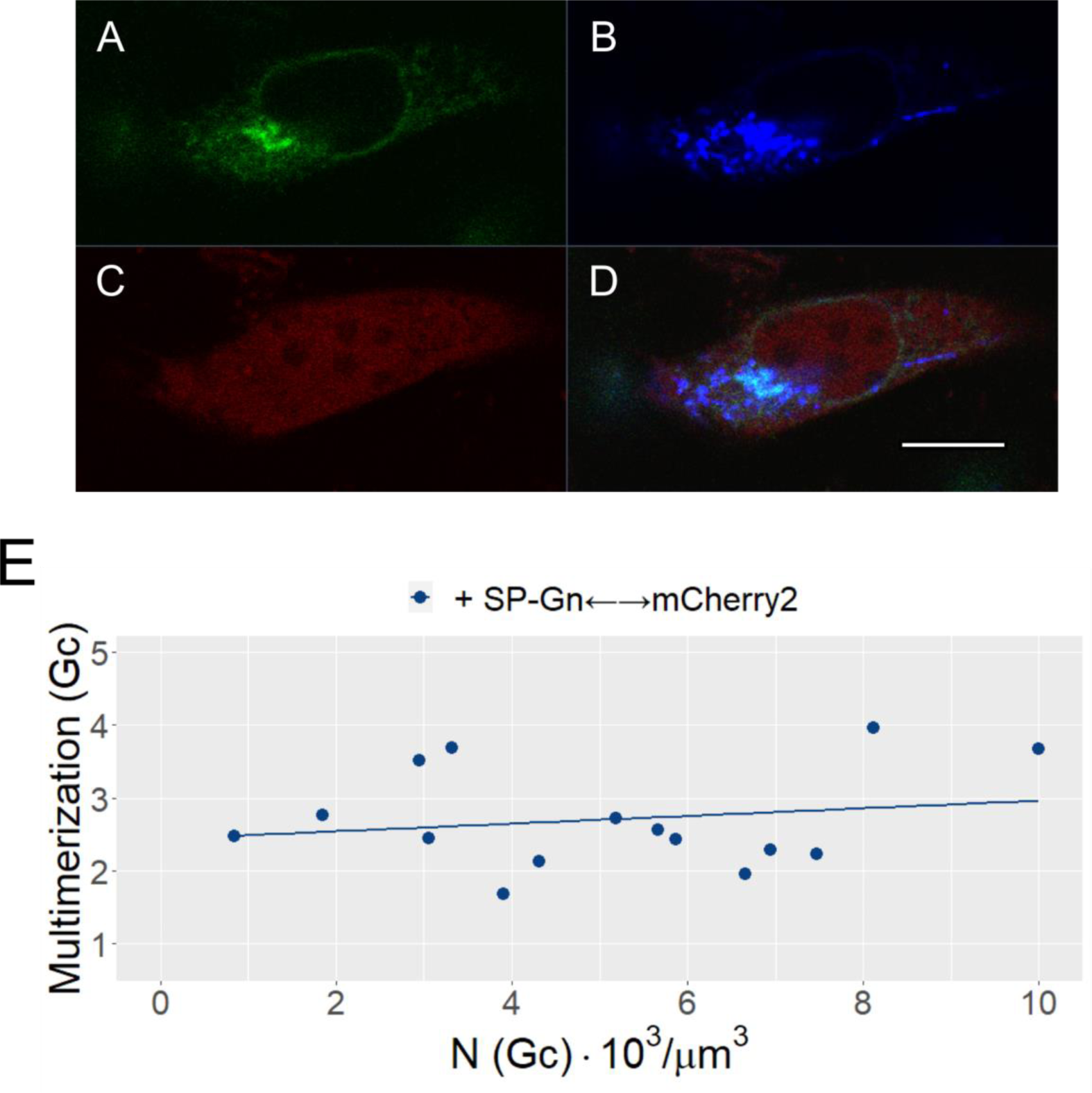
Concentration-dependent oligomerization of PUUV Gc in the presence of Gn. CHO-K1 cells expressed SP-mEGFP-Gc and SP-Gn (together with cytosolic mCherry2 within a bi-directional vector; indicated as SP-Gn⟵⟶mCherry2). A-D) Representative confocal microscopy image of CHO-K1 transfected with SP-mEGFP-Gc and SP-Gn⟵⟶mCherry2 vectors. Split view of SP-mEGFP-Gc (A), Golgi-mTurquoise marker (B), cytosolic mCherry2 (C) and merged channels (D). White bars are 5 μm. B) SP-mEGFP-Gc multimerization as a function of total Gc concentration (in monomer units), obtained via N&B analysis as described in the Materials and Methods section. The analysis was restricted to cells which showed significant expression of mCherry2. Each point is calculated from the average brightness extracted from ROIs in 2-3 cells with similar protein concentrations. Solid lines are a fit to an empirical model (see Materials and Methods) and are meant as guide to the eye. Each dataset is obtained from at least 4 separate experiments, for a total of ca. 35 cells.

## Discussion

The formation of homo- and hetero-complexes between Gn and Gc proteins is supposed to drive the assembly of new virions in infected cells. In spite of the importance of these inter-molecular interactions, there is a general lack of information regarding the oligomeric state of HV GPs directly during the process of virus assembly. In this study, we addressed this question and quantified the interactions that lead to oligomeric spike complex formation in the physiological context of living cells.

### Limited Gn and Gc homotypic-interactions

In the case of Gc singularly expressed in different cell lines, the protein was found in monomeric and dimeric form both in the ER and GA (Fig. 1 D, 3 C, S3, S4). On the other hand, Gn oligomers ranged in size between monomers and tetramers, with a considerable correlation to the local protein concentration in the ER (Fig. 1 B). These results validate the hypothesis that HV spike formation is modulated by local protein concentration (8). Also, the formation of Gn tetramers and dimeric Gc:Gc contacts is in agreement with the proposed arrangement of GPs on the surface of PUUV and TULV particles, consisting of Gn-Gc hetero-octamers (i.e. four Gn and four Gc) interconnected via a small Gc:Gc interaction patch (18, 20, 38). By using mutations that stabilize (H303C) or destabilize (H303E) the Gc:Gc contact region between spikes on viral particles (23), we observed by N&B that the Gc-Gc interactions that occur in the GA of living cells are directed by the same contact interface in absence of Gn. The H303C Gc substitution induced dimers very efficiently while, conversely, the H303E substitution hindered dimer formation (Fig. 3). We were thus able to discern subtle changes in the apparent monomer-dimer equilibrium directly in living cells and validated previous results that identified specific amino acids at the Gc:Gc inter-spike contact interface as sensitive mediators of this interaction. The Gc:Gc contact surface might be too small to allow the formation of stable Gc dimers in solution, according to the observation that soluble Gc ectodomain remains monomeric in solution (22, 39). However, our results suggest that the confinement of Gc in a two-dimensional geometry within lipid membranes in the GA (and the consequent high local protein concentration) are sufficient to allow Gc:Gc association in living cells. These interactions might be further favored by inter-spike restrains during viral assembly and on the viral surface.

Similarly, we also analyzed regions in Gn which are responsible for its quaternary structure. We tested the concentration-dependent multimerization of the mutant ΔGn in which the N-terminal, membrane distal lobes of Gn (21) were deleted (while maintaining the Gn central stalk, transmembrane domain and endodomain). The fact that this Gn mutant also showed increasing multimerization from monomers to tetramers – correlating to the local protein concentration (Fig. 4) – suggests that the membrane-distal lobe of Gn is not needed for Gn multimerization. The ca. 4-fold higher concentrations that were required for a significant multimerization to occur indicate that the membrane-distal lobe might contribute to the tetramer stabilization. The lower stability of ΔGn tetramers might also be partially caused by the N-terminal fusion FPs which, despite the 14 amino acid long linker region, may produce partial hindrance at the tetrameric center of the ectodomain, making multimerization less efficient. We previously observed an analogous behavior also for the Influenza A GP hemagglutinin, whose TMD can multimerize although only to a limited extent, compared to the wild-type (30).

### Multimerization-concentration curves

In previous investigations from our laboratory, we have discussed the possibility of Gn tetramerization in cell models exclusively expressing this viral GP (16). Nevertheless, in that case, only average multimerization values could be obtained and the presence of tetramers had to be assumed *a priori* in order to estimate the amounts of the different oligomeric species. In this extended investigation instead, we have explored a wide range of concentrations and quantified GP multimerization with single-cell resolution. Further, the observation that the apparent multimeric state of Gn saturates at a value of ∼4 (Fig. 1 B) proves that Gn tetramers are indeed present (being the predominant multimeric species at concentrations higher than ca. 2000 proteins /µm^3^) and the amount of larger complexes is negligible at all concentrations. A similar conclusion can be drawn about Gc multimerization approaching asymptotically a value of ∼2, indicating the presence of a monomer-dimer equilibrium, without the need for *a priori* assumptions deriving from biochemical/structural data.

### GP multimerization is consistent for different viral strains and cellular environments

Comparison of different HV strains suggests only minor differences in behavior between PUUV and HTNV GPs expressed in CHO-K1 cells (Fig. 2). This is in agreement with the previous observation that, despite the genetic and geographical distinction between strains, the molecular structure of HV GPs are highly conserved (21, 22, 38-40). More in detail, while Gn localizes in the ER for both strains, Gc showed a partial enrichment in the GA for PUUV but not for HTNV, in agreement with a previous report (14). In this context, it is worth noting that former investigations provided conflicting results regarding the intracellular localization of singularly expressed Gn and Gc (12-14). Here, we report a consistent multimerization behavior for both Gn and Gc, when expressed alone, not correlating with their intracellular localization or cell type. For example, we observed that Gc is found in monomeric or dimeric form, independently from viral strain and localization in GA/ER (Fig. 2, S3).

### Gn-Gc large scale hetero-interactions in the GA

While the investigation of singularly expressed GPs can already provide information about the formation of homotypic complexes, it must be noted that HV assembly in physiological conditions occurs exclusively in the presence of both Gn and Gc. The last part of our study deals therefore with the quantification of protein-protein interactions occurring in the presence of both GPs. In detail, it was shown for HTNV (13, 14), PUUV (16), ANDV and SNV (15, 17) that Gn and Gc expressed together (from a single GPC, separate cDNAs or in the context of infection) co-localize in the GA. However, in this study, we quantify for the first time the oligomerization states of each protein in connection to its subcellular localization, its concentration and the concentration of its interaction partner. First, we confirmed that both PUUV Gn and Gc tagged with FPs are enriched in the GA. In this case, though, Gn does not form large oligomers and shows even decreased multimerization extent than when expressed alone (Fig. 5 E). We believe that the decreased oligomerization capabilities of Gn in this case are probably due to the steric hindrance generated by the fluorescent tag present on Gc N-terminus, since SP-mEGFP-Gn alone can readily form tetramers. This result, while originating from a drawback commonly encountered in fluorescence microscopy investigations, has some general implications. First, this is a strong indication of a direct Gn-Gc interaction, since a part of the Gc fluorescent construct appears to interfere with Gn-Gn contacts. Second, the formation of even relatively small Gn-Gc hetero-oligomers (Gn remaining monomeric or dimeric in average) seems sufficient for Gn transport to the GA. Third, it demonstrates that Gn-Gc interactions do not require the previous formation of Gn homo-tetramers (in disagreement with assembly model 1 proposed by Hepojoki et al. (24)).

The possibility that Gn-Gc multimerization is hindered when both GPs contain a fluorescent label is confirmed by the observation that, in the presence of unlabeled Gc, Gn forms multimers with up to eight subunits in average. We therefore developed a novel quantitative fluorescence approach to monitor the interactions between a fluorescent protein and an unlabeled protein, while also being able to estimate the concentrations of both interacting partners (see Figs. 5 E and 6). Using this approach, we observed that, in the presence of non-fluorescent Gc, Gn forms large oligomers, containing up to 8-12 subunits. It must be noted that, in contrast to the case of singularly expressed Gn, we did not observe saturation of Gn multimerization even for the highest concentrations considered. This suggests that even larger Gn multimers might be present in this samples and the observed multimerization is in fact an average estimate. Also, the weak correlation between Gn multimerization and Gc concentration (Fig. S8) indicates that the formation of larger Gn complexes does not require preformed Gc:Gc dimeric association (that occurs at high Gc concentrations, see Fig. 1 D). Instead, the results support the idea that Gn and Gc may associate already at low concentrations, subsequently allowing the formation of higher order Gn-Gc species.

For the case of Gc in the presence of (non-fluorescent) Gn, instead, the apparent multimerization ranges between two and four, in average (i.e. oligomers with more than four units might also be present, albeit in small amounts). Since only Gc dimers are observed at best in the absence of Gn, multimers containing more than two Gc units must be mediated by Gn oligomers. The same holds true for Gn tetramers as largest Gn oligomer in the absence of Gc (i.e. four Gn units). The observation of Gc complexes with approximately four units diffusing as a whole might be explained by several Gn-Gc configurations, such as those shown in Fig. 7 A-D.

**Figure 7:**
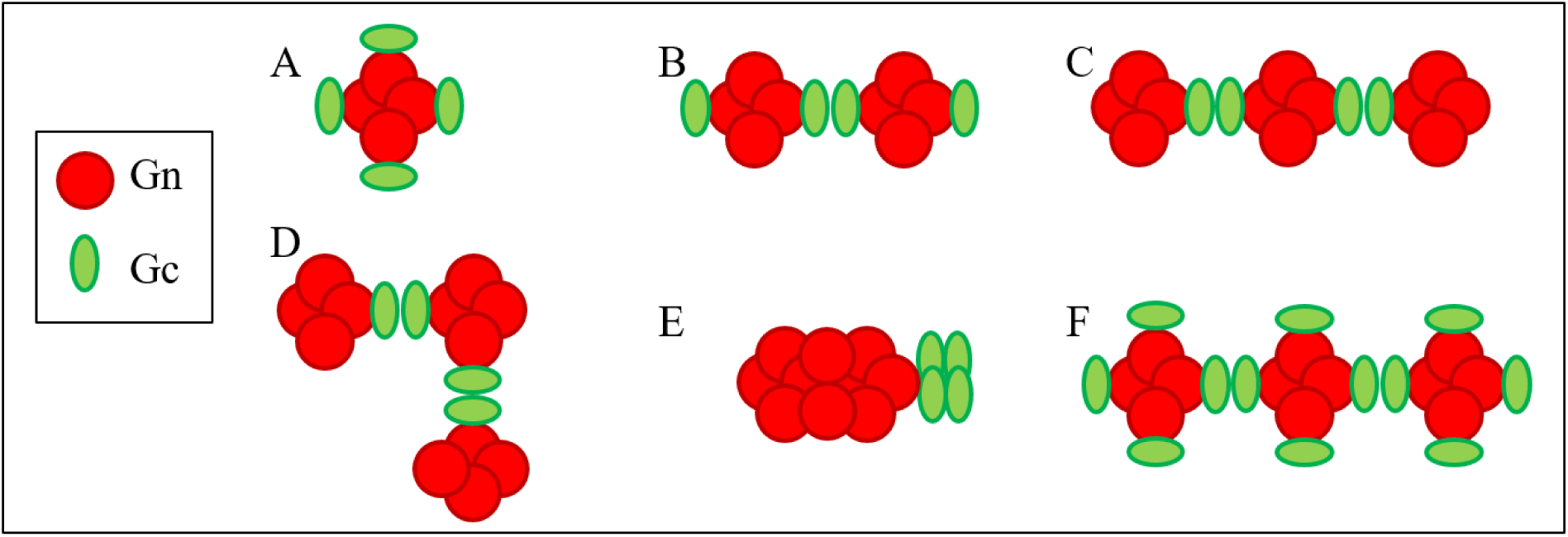
Examples of Gn-Gc hetero-multimers compatible and incompatible with the highest observed multimerization states in cells expressing both Gn and Gc. A-D) Examples of hetero-multimers containing 4 Gc units and 4-12 Gn units. E) Gn-Gc arrangement containing a Gn homo-decamer linked to a Gc tetramer. Such hetero-multimer is compatible with the results shown in Figs. 5 and 6, but not with the GP quaternary structure observed for Gn (or Gc) in the absence of the other GP (Fig. 1). F) Gn-Gc hetero-octamers connected via Gc:Gc contacts described for the surface of viral particles (21, 23).

Larger Gn-Gc hetero-multimers, such as those described in Fig. 7 B-D, are additionally in line with the observation of high multimeric states for Gn mentioned before. Also, alternative spatial arrangements of Gn-Gc hetero-multimers (see e.g. the large Gn homo-multimer linked to a Gc homo-tetramers in Fig. 7 E) would not be compatible with the largest Gn (or Gc) homo-oligomers observed in the absence of the other GP, i.e. tetramers (or dimers).

Clearly, we cannot exclude that the limited degree of Gc oligomerization (i.e. the limited amount of Gc monomers participating in the described possible Gn-Gc hetero-multimers) might be partially due to the presence of the N-terminal fluorescent tags. Our experiments cannot rule out the possibility that, in the absence of FPs, more Gc monomers might be present in Gn-Gc hetero-multimers (see e.g. Fig. 7 F). It is important to stress therefore that the FPs might have prevented a Gn-Gc stoichiometry of 1:1 that is otherwise guaranteed in natural infections. Nevertheless, it must be noted these results validate in general the proposed GP organization on the viral surface as Gn tetramers connected via Gc:Gc contacts (21, 23, 24). Finally, the lack of correlation between Gc multimerization and Gn concentration suggests once more that the assembly model 1 (24) cannot be entirely accurate, since that would require the formation of Gn multimers (i.e. higher Gn concentrations) for Gc multimerization to occur.

## Conclusions

We have investigated the multimerization of HV envelope protein Gn and Gc from “old-world” HV, both singularly or expressed together in living cells, using quantitative fluorescence microscopy approaches. Based on our findings, we can validate a spike assembly model (based on the assembly model 2 proposed by Hepojoki et al. (24)) according to which Gn-Gc small hetero-oligomers (formed e.g. by a Gn dimer and a Gc monomer or, simply, a Gn-Gc hetero-dimer) are first formed in the ER allowing translocation of this complex to the GA. There, Gn-Gn interactions drive the formation of the hetero-octameric complex and Gc stabilizes contacts between neighboring spikes. The further assembly into still larger complexes is reasonably hindered by the presence of FPs in this case. While the presented method appears thus inherently limited by the influence of the fluorescent labels, it is worth stressing that we provide here, for the first time, direct evidence for the initial steps of HV assembly in the GA of living cells. The non-disruptive method presented in this work opens therefore the possibility for further investigations in the context of HV assembly, including for example the characterization of the interaction between GPs and the ribonucleocapsid protein (RNP) or other viral components.

## Acknowledgements

This work was supported by the German Research Foundation (DFG grant 407961559 to S.C.) and ANID (Chile) grants FONDECYT 1181799 and Programa de Apoyo a Centros con Financiamiento Basal 170004 to N.D.T. The authors thank V. Dunsing for useful discussion.

## Contributions

Research planning, N.D.T. and S.C.; investigation, R.A.P. and A.A.K.; data analysis, R.A.P. and A.A.K.; writing–original draft preparation, R.A.P. and S.C.; writing–review and editing, A.A.K. and N.T.; software, R.A.P.; supervision, S.C.; funding acquisition, N.D.T. and S.C.

