## Supplementary Info for "Detection of Envelope Glycoprotein Assembly from Old-World Hantaviruses in the Golgi Apparatus of Living Cells"

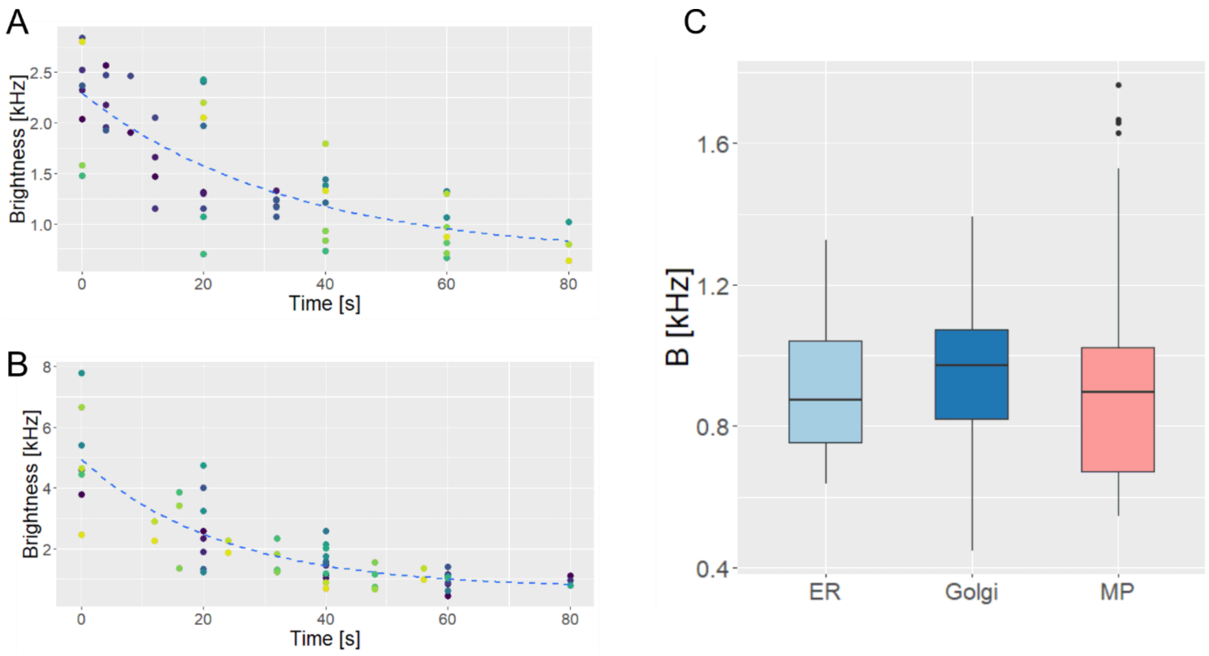


**Figure S1: Brightness-bleaching curves of ER-mEGFP (A) and Golgi-mEGFP (B)**. Absolute brightness was measured at different time points, while repeatedly bleaching the sample using high excitation powers (see Materials and Methods). Each color represents one series of measurements performed on a single cell. Dashed lines show an exponential decay fitted to the experimental data, as guide to the eye. The minimum values (corresponding to the flattening of the curve) for the measured brightness should correspond to that of monomers, both for ER-mEGFP (A) and Golgi-mEGFP (B). Asymptotic values are represented as box plots (C), together with the brightness of the standard monomer reference for the plasma membrane (MyrPalm-mEGFP), measured directly (i.e. without bleaching cycles). The data for ER-mEGFP and Golgi-mEGFP were extracted from the bleaching curves of 15-20 cells from 3 separate experiments. The data for MyrPalm-mEGFP was collected from 38 cells in 4 different experiments.

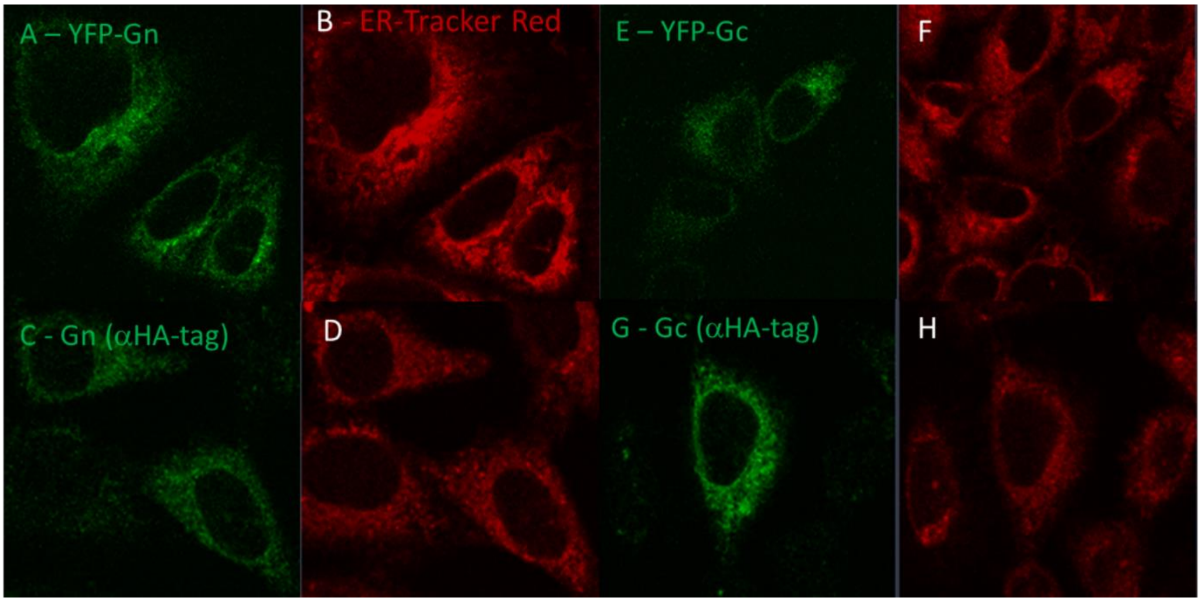


**Figure S2: Gn and Gc intra-cellular localization.** A-B) Representative confocal fluorescence images of CHO cells expressing SP-mEYFP-Gn (A) and a mCherry-labelled ER marker (B). C-D) Representative confocal fluorescence images of CHO cells expressing HAtag-Gn labelled with an anti-HAtag rabbit primary antibody and an Alexa488-labelled secondary antibody (C) and a mCherry-labelled ER marker (D). E-F) Representative confocal fluorescence images of CHO cells expressing SP-mEYFP-Gc (E) and a mCherry-labelled ER marker (F). G-H) Representative confocal fluorescence imaging of CHO cells expressing HAtag-Gc labelled with an anti-HAtag rabbit primary antibody and an Alexa488-labelled secondary antibody (G) and a mCherry-labelled ER marker (H). All imaging was performed at room temperature. Image size is 40 µm x 40 µm (50 µm x 50 µm for panels E-F).

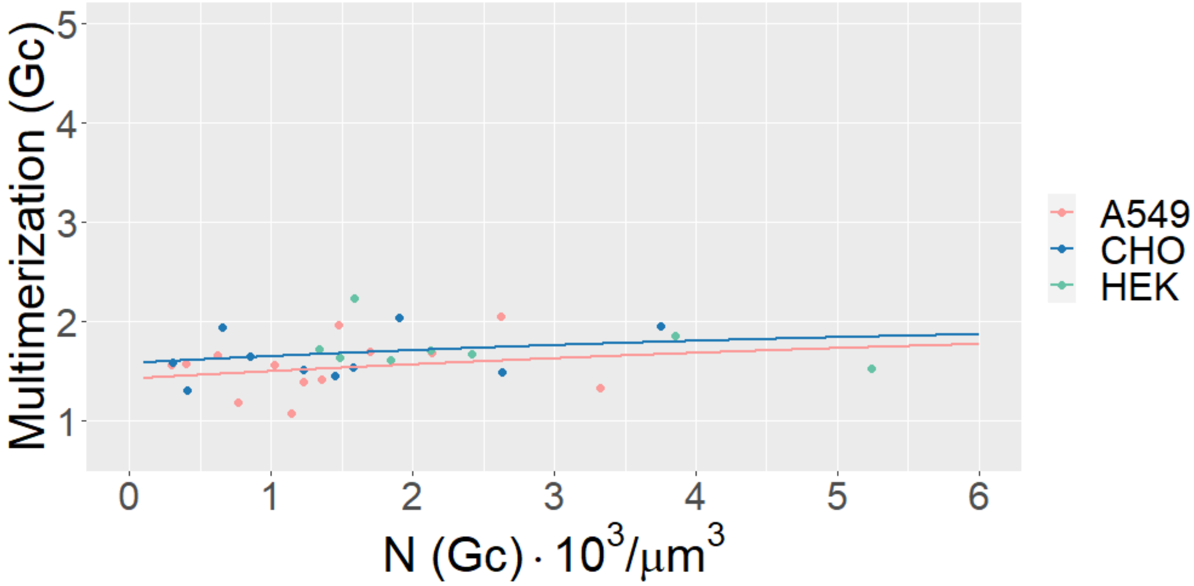


**Figure S3: Concentration-dependent oligomerization of Gc in the ER.** Different cell lines were transfected with SP-mEGFP-Gc. N&B analysis was applied to a ROI in each cell identified by an ER-mCherry marker. The graph shows SP-mEGFP-Gc multimerization as a function of its total monomer concentration. Multimerization values were obtained from molecular brightness normalization, as described in the Materials and Methods section. Each point is calculated from the average brightness extracted from ROIs in 2-4 cells with similar protein concentrations. Solid lines are fit to an empirical model (see Materials and Methods) and are meant as guide to the eye. Each dataset is obtained from at least 3 independent experiments, for a total of ca. 30-50 examined cells.

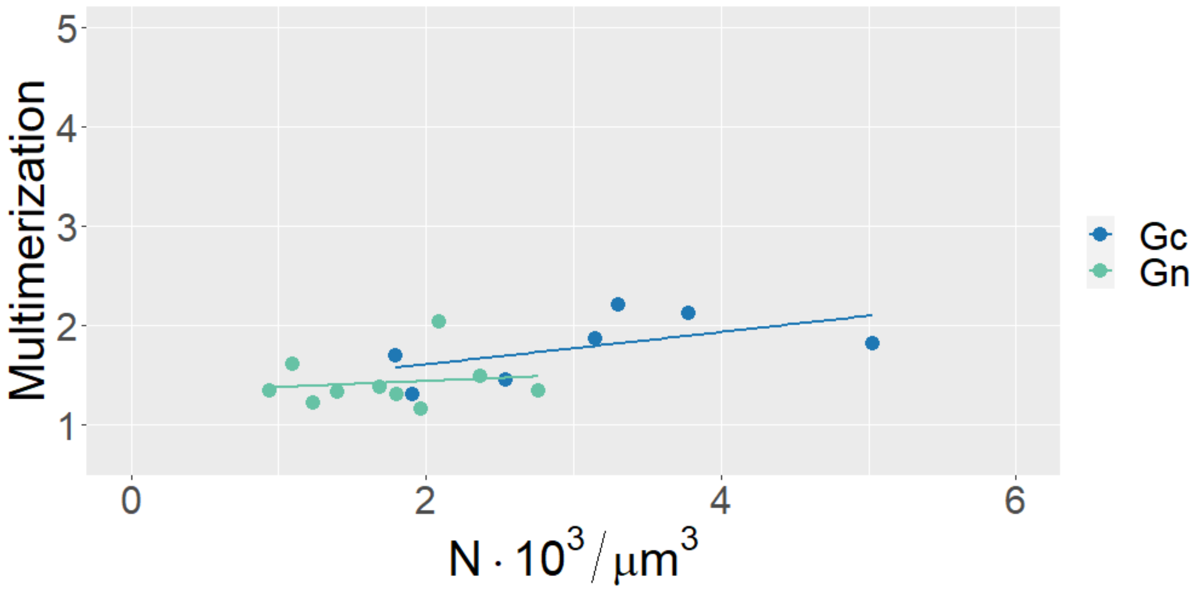


**Figure S4: GP multimerization as a function of total monomer concentration in VeroE6 cells.** VeroE6 cells were transfected with either SP-mEGFP-Gn or SP-mEGFP-Gc. N&B analysis was applied to a ROI in each cell identified by ER-mCherry or GA-mCherry markers. The graph shows GP multimerization as a function of total monomer concentration. Multimerization values were obtained from molecular brightness normalization, as described in the Materials and Methods section. Each point is calculated from the average brightness extracted from ROIs in 2-4 cells with similar protein concentrations. Solid lines are guide to the eye. Each dataset is obtained from at least 3 independent experiments, for a total of ca. 25-35 examined cells.

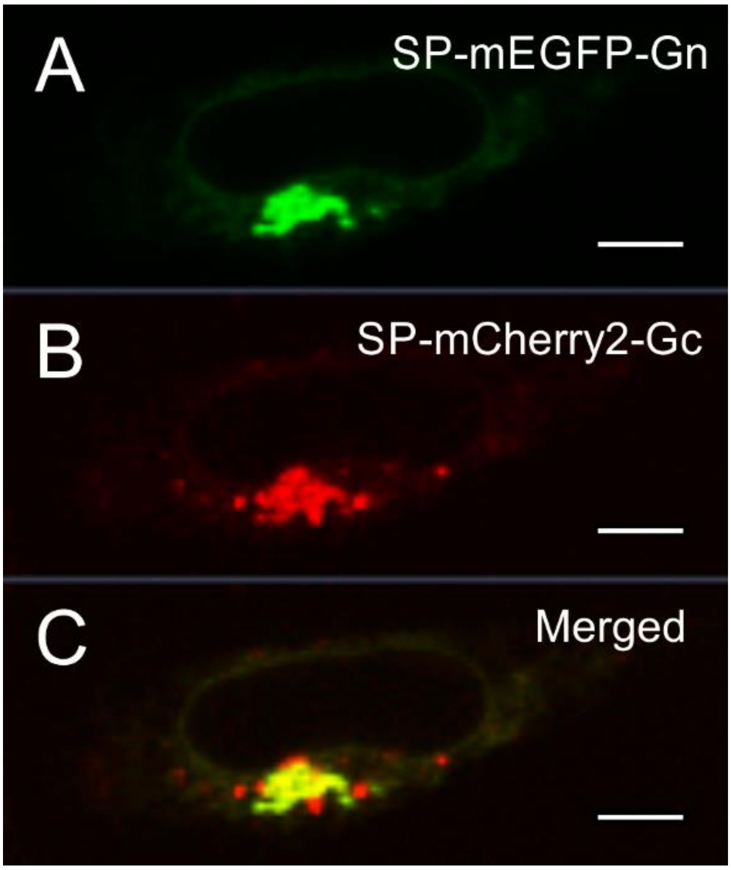


**Figure S5: Intracellular distribution of SP-mEGFP-Gn and SP-mCherry2-Gy upon co-expression.** Representative confocal fluorescence images of CHO cells expressing SP-mEGFP-Gn (A) and SP-mmCherry2-Gc (B). Scale bars are 5 µm.

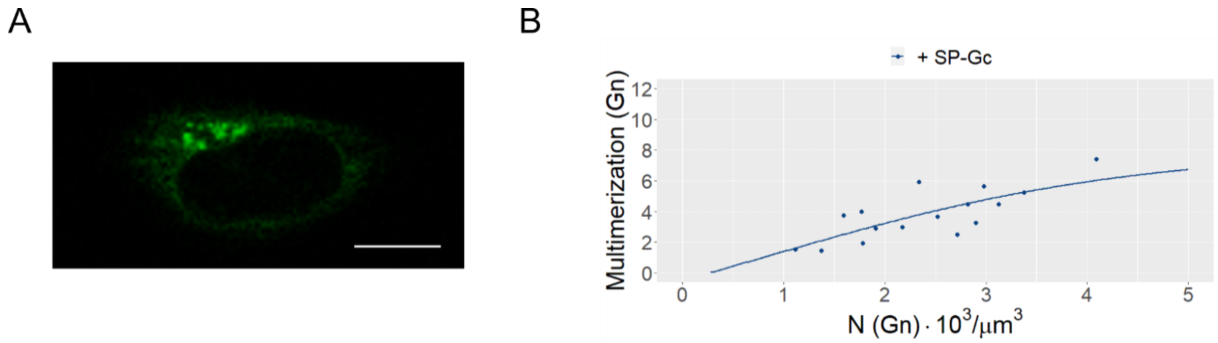


**Figure S6: Concentration-dependent oligomerization of PUUV Gn in the presence of Gc.** CHO-K1 cells were co-transfected with SP-mEGFP-Gn and SP-Gc. A) Representative confocal microscopy image of CHO-K1 transfected with SP-mEGFP-Gn and SP-Gc. Scale bar is 5 µm. The presence of both GPs is inferred by the non-homogeneous distribution of SP-mEGFP-Gn between ER and GA. The intracellular distribution pattern of the protein in this case is comparable e.g. to that observed in the presence of both GPs (e.g. Fig. 5 in the main text or Fig. 3 in (1)). B) N&B analysis of SP-mEGFP-Gn multimerization was performed as above mentioned and restricted to cells which showed significant localization of SP-mEGFP-Gn in the GA region. The graph shows SP-mEGFP-Gn multimerization as a function of its total monomer concentration. Each point is calculated from the average brightness extracted from ROIs in 2-3 cells. Solid lines are a fit to an empirical model (see Materials and Methods) and are meant as guide to the eye. Each dataset is obtained from at least 4 separate experiments, for a total of ca. 30 cells.

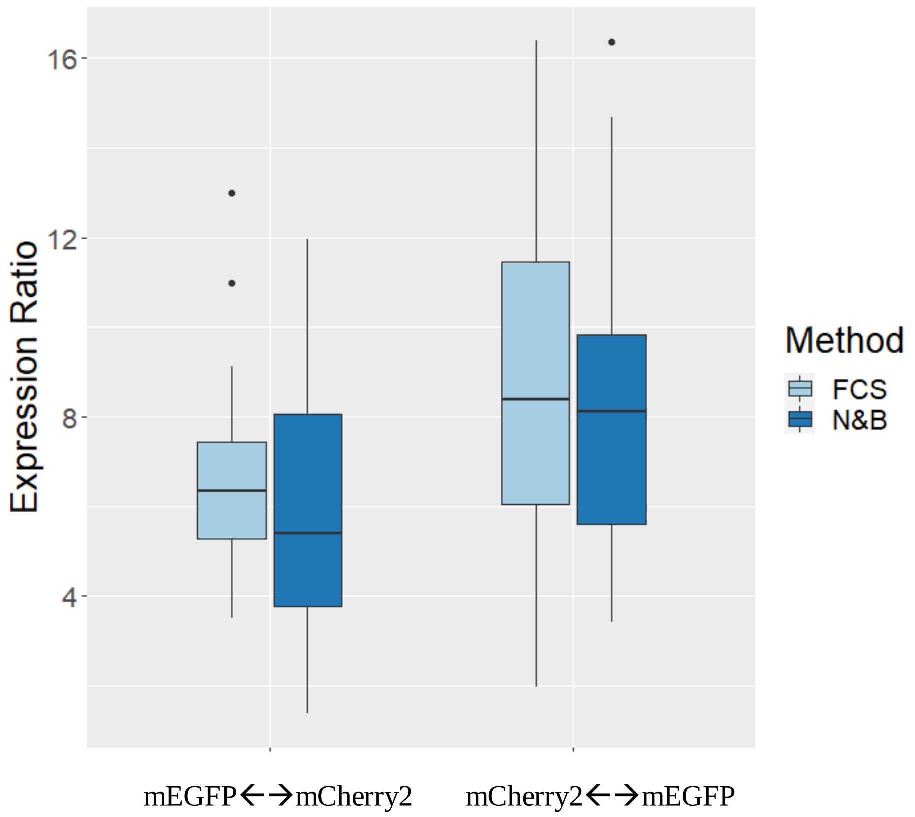


**Figure S7: Calibration of bi-directional plasmids.** We examined bidirectional plasmids with either i) mEGFP upstream and mCherry2 downstream of the bidirectional promoter region (mEGFP←→mCherry2) or ii) mCherry2 upstream and mEGFP downstream of the bidirectional promoter region (mCherry2←→mEGFP). Point FCS and N&B measurements were independently performed in the cytosol of transfected CHO-K1 cells to calculate the concentrations of each FP (which appeared roughly homogeneously distributed in the whole cell), for each bidirectional plasmid. The box plot shows the apparent "expression ratio", defined as the ratio between the measured amounts of FPs (downstream und upstream of the promoter region, respectively). Each FP amount is quantified as the number of molecules detected in the confocal volume, provided by FCS and N&B. The difference between the two ratios can be qualitatively explained by the slightly lower fluorescence probability $p_m$ of mCherry2 (ca. 0.62), compared to mEGFP (ca. 0.72) (2). Assuming in fact that the actual expression ratio does not depend on the identity of the expressed protein, the expression ratio measured with a fluorescence-based method for the vector mCherry2←→mEGFP would additionally contain a multiplicative factor $\frac{p_m(\text{mCherry2})}{p_m(\text{mEGFP})}$. For the vector mEGFP←→mCherry2, the multiplicative factor would be $\frac{p_m(\text{mEGFP})}{p_m(\text{mCherry2})}$.

The expression ratio for mEGFP←→mCherry2 was then further used to estimate the concentration of Gc or Gn for the experiments in which the SP-Gc←→mCherry2 or SP-Gn←→mCherry2 constructs were used for transfection.

To calculate GP concentrations in the GA, we first measured the number of soluble mCherry2 proteins detectable in the confocal volume ($N_{mCh2}$) from point FCS measurements. Next, we calculated $N_{GP-cell}$, i.e. the corresponding number of GPs in the confocal volume, in the case that the protein had the same intra-cellular distribution of mCherry2. $N_{GP-cell}$ is also simply connected to the total number of GP monomers in the whole cell $N_{GP}^{total}$ :

$$N_{GP-cell} = \frac{N_{GP}^{total}}{V_{cell}} = \frac{N_{mCh2}}{\text{Expression Ratio from mEGFP[mCherry2]}},$$

where $V_{cell}$ is the total cell volume in confocal volume units. For the sake of simplicity, we assumed here that the probability of obtaining a stable, correctly formed GP is similar to the probability of obtaining a correctly folded, detectable mEGFP (i.e. ca. 70-75%).

Next, we estimated a GA-partition coefficient K (for either Gn or Gc), defined as:

$$K = \frac{(N_{GA} \cdot A_{GA})}{(N_{GA} \cdot A_{GA} + N_{ER} \cdot A_{ER})},$$

where $N_{GA}$ is the average number of GPs in a volume unit within the GA, $A_{GA}$ is the apparent surface area occupied by the GA in the examined confocal image (e.g. in pixel units), $N_{ER}$ is the average number of GP in a pixel belonging to ER and $A_{ER}$ is the apparent surface area occupied by the ER in the examined confocal image. K was obtained as average value from the analysis of 15-20 cells expressing either Gn or Gc respectively. We assume here that the ER and the GA have roughly the same depth and that the GPs partition exclusively in the ER and GA.

The estimated number of the GP monomers in a confocal volume position in the GA is then:

$$N\text{ (Gn or Gc)} = \frac{N_{GP-cell} \cdot K \cdot V_{cell}}{V_{GA}},$$

where $V_{GA}$ is the volume of the GA in confocal volume units. $V_{GA}/V_{cell}$ was assumed to be ~0.04 (3). For graphical representation in Figs. S8 and S9, N(Gn or Gc) was finally converted in $\mu m^{-3}$ units.

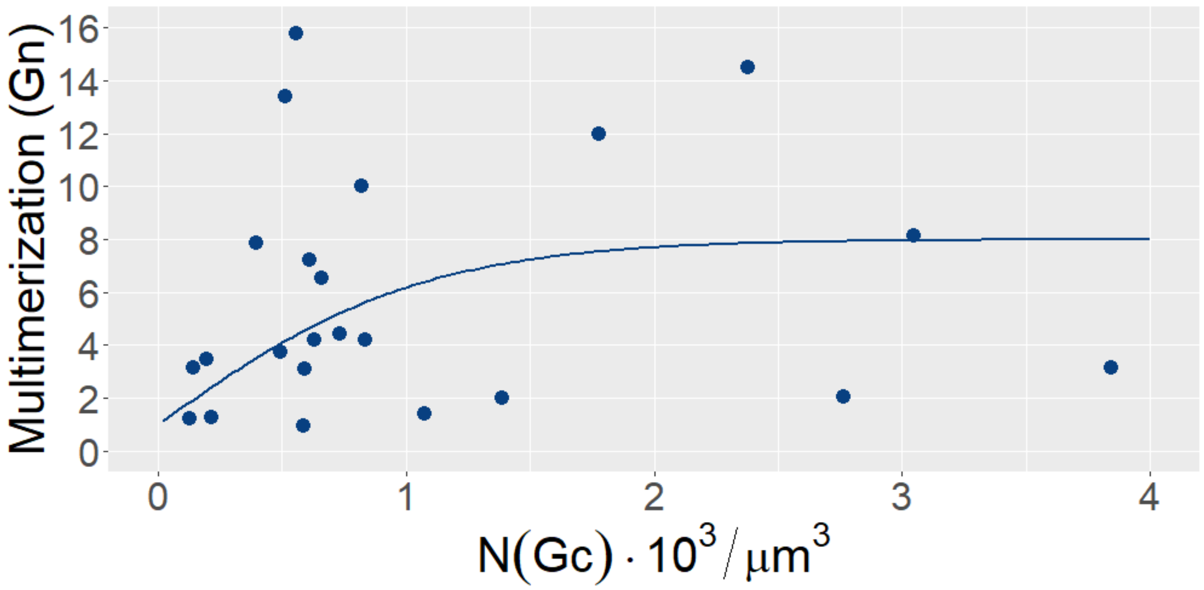


**Figure S8: Gn multimerization as a function of Gc total monomer concentration.** CHO-K1 cells were co-transfected with SP-mEGFP-Gn and SP-Gc←→mCherry2. N&B analysis of SP-mEGFP-Gn multimerization was performed as above mentioned and restricted to cells which showed significant expression of mCherry2. Multimerization of SP-mEGFP-Gn is expressed as a function of the estimated SP-Gc concentration in the GA, calculated from FCS measurements of the cytosolic mCherry2 (see Fig. S7). Each point is calculated from the average brightness extracted from ROIs in one cell. Solid lines are a fit to an empirical model (see Materials and Methods) and are meant as guide to the eye. Each dataset is obtained from at least 4 separate experiments, for a total of 24 cells.

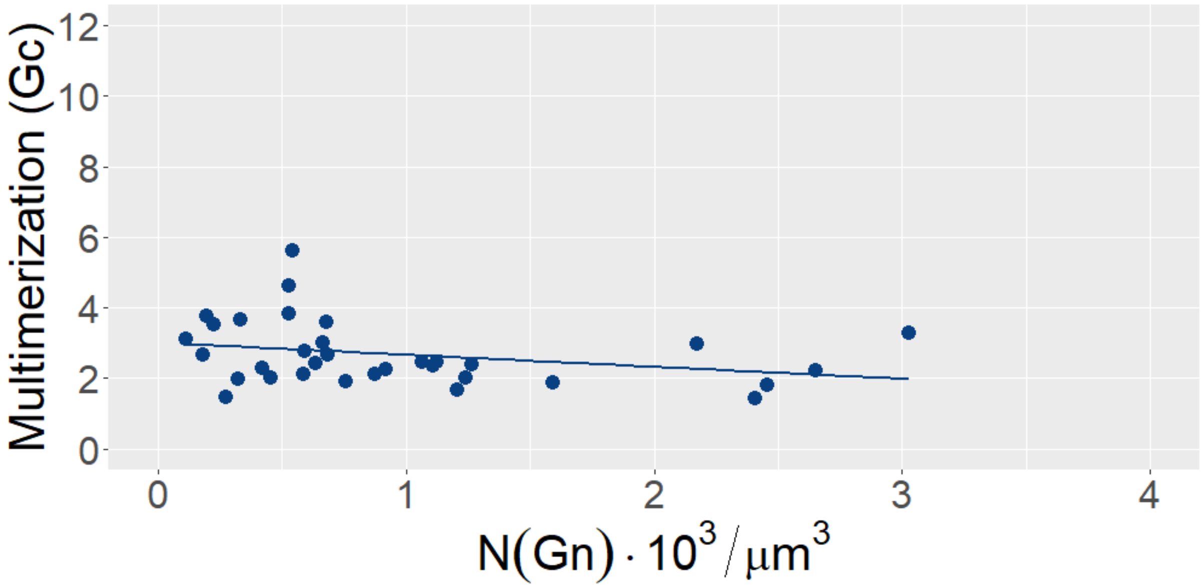


**Figure S9: Gc multimerization as a function of Gn total monomer concentration.** CHO-K1 cells were co-transfected with SP-mEGFP-Gc and SP-Gn←→mCherry2. N&B analysis of SP-mEGFP-Gc multimerization was performed as above mentioned and restricted to cells which showed significant expression of mCherry2. Multimerization of SP-mEGFP-Gc is expressed as a function of the estimated SP-Gn concentration in the GA, calculated from FCS measurements of the cytosolic mCherry2 (see Fig. S7). Each point is calculated from the average brightness extracted from ROIs in one cell. Solid lines are a fit to an empirical model (see Materials and Methods) and are meant as guide to the eye. Each dataset is obtained from at least 4 separate experiments, for a total of 33 cells.

1. **Sperber HS, Welke RW, Petazzi RA, Bergmann R, Schade M, Shai Y, Chiantia S, Herrmann A, Schwarzer R.** 2019. Self-association and subcellular localization of Puumala hantavirus envelope proteins. Sci Rep **9:**707.
2. **Dunsing V, Luckner M, Zuhlke B, Petazzi RA, Herrmann A, Chiantia S.** 2018. Optimal fluorescent protein tags for quantifying protein oligomerization in living cells. Sci Rep **8:**10634.
3. **Sturgess JM, De la Iglesia FA.** 1972. Morphometry of the golgi apparatus in developing liver. J Cell Biol **55:**524-530.
